# A deep learning-based model of normal histology

**DOI:** 10.1101/838417

**Authors:** Tobias Sing, Holger Hoefling, Imtiaz Hossain, Julie Boisclair, Arno Doelemeyer, Thierry Flandre, Alessandro Piaia, Vincent Romanet, Gianluca Santarossa, Chandrassegar Saravanan, Esther Sutter, Oliver Turner, Kuno Wuersch, Pierre Moulin

## Abstract

Deep learning models have been applied on various tissues in order to recognize malignancies. However, these models focus on relatively narrow tissue context or well-defined pathologies. Here, instead of focusing on pathologies, we introduce models characterizing the diversity of normal tissues. We obtained 1,690 slides with rat tissue samples from the control groups of six preclinical toxicology studies, on which tissue regions were outlined and annotated by pathologists into 46 different tissue classes. From these annotated regions, we sampled small patches of 224 × 224 pixels at six different levels of magnification. Using four studies as training set and two studies as test set, we trained VGG-16, ResNet-50, and Inception-v3 networks separately at each of these magnification levels. Among these models, Inception-v3 consistently outperformed the other networks and attained accuracies up to 83.4% (top-3 accuracy: 96.3%). Further analysis showed that most tissue confusions occurred within clusters of histologically similar tissues. Investigation of the embedding layer using the UMAP method revealed not only pronounced clusters corresponding to the individual tissues, but also subclusters corresponding to histologically meaningful structures that had neither been annotated nor trained for. This suggests that the histological representation learned by the normal histology network could also be used to flag abnormal tissue as outliers in the embedding space without a need to explicitly train for specific types of abnormalities. Finally, we found that models trained on rat tissues can be used on non-human primate and minipig tissues with minimal retraining.

**Author contribution:** T.S. and H.H. contributed equally to this work.

**Significance statement:** Like many other scientific disciplines, histopathology has been profoundly impacted by recent advances in machine learning with deep neural networks. In this field, most deep learning models reported in the literature are trained on pathologies in specific tissues/contexts. Here, we aim to establish a model of normal tissues as a foundation for future models of histopathology. We build models that are specific to histopathology images and we show that their embeddings are better feature vectors for describing the underlying images than those of off-the shelf CNN models. Therefore, our models could be useful for transfer learning to improve the accuracy of other histopathology models.

## Introduction

After breakthrough results in the ImageNet challenge 2012 (Krizhevsky et al., 2012) and subsequent challenges, deep neural networks have led to a radical transformation of many scientific disciplines, including medicine. In particular, deep learning-based diagnostics have achieved physician-level accuracy across a wide range of diagnostic tasks (Esteva et al., 2019; Liu et al., 2019), and the potential of deep learning to transform health care has been broadly recognized by regulatory agencies as well as in the literature (Hinton, 2018; Naylor, 2018).

The deep learning transformation of computational pathology (Litjens et al., 2017) faces specific challenges posed by whole-slide imaging (WSI) of tissue sections stained with hematoxylin and eosin (H&E) including large image sizes, color variation and other artefacts, and more importantly the multiscale nature of the data – with various emergent structures from the nuclei to the organ level (Komura and Ishikawa, 2018). Moreover, obtaining annotations is more challenging for histopathology than other disciplines: it is highly time-consuming and requires input from trained pathologists.

To date, the bulk of machine learning approaches to digital pathology have focused on detection and segmentation of histologic primitives such as nuclear size and shape, grading of lesions (Campanella et al., 2019), prediction of clinical outcomes, and linking histopathology with other data types (Komura and Ishikawa, 2018; Madabhushi and Lee, 2016). Recent community challenges aimed at detecting lymph node metastases (CAMELYON16: Bandi et al., 2019; CAMELYON 17: Ehteshami Bejnordi et al., 2017), assessing tumor proliferation in breast cancer (TUPAC16: Veta et al., 2019), or detecting and classifying lung cancer (ACDC-LungHP, 2019). Other recent work has explored the prediction of genetic alterations from H&E-stained slides (Coudray et al., 2018; Kather et al., 2019).

All these applications operate in a disease-specific context, which presupposes a histopathological diagnosis to be established prior to submitting slides to the application. That is, the morphological features learned by such models are optimized for the differentiation of particular outcomes given the initial diagnosis. Therefore, it is difficult to imagine how such models could contribute to applications aiming at assisting the diagnosis, i.e. identify key diagnostic features from a non-predetermined histopathological slide. On the contrary, a model based on features that differentiate all possible tissue characteristics pertaining to identifying any type of tissue or histological lesion would provide invaluable support for the development of applications assisting pathological diagnosis. The mere automated identification of samples devoid of lesion could tremendously improve the efficiency and quality of pathology evaluation in areas of clinical practice providing screening for lesions, but also in preclinical development of medicines, where pathology is pivotal for the assessment of drug safety.

Our work focused on the development of a holistic model of histology, which, from our perspective, will lay the foundation of more complex models suitable for assistance to diagnosis. Obtaining a complete and homogeneous set of normal human tissues through autopsies is hindered by several factors, including the necessary consents, the age of the patients, or the delays between death and collection. In contrast, preclinical toxicology studies comprise control animals which are not exposed to chemical compounds and where all organs are systematically collected. This immense collection of histological slides of normal animal tissues represents a significant opportunity to develop holistic tissue models. Moreover, the histological structure of the tissues is largely conserved between the species used in drug development and humans.

The three key contributions of this article are the following:

### Tissue recognition

First, we show that a comprehensive set of mammalian tissues can be recognized by standard convolutional neural networks (CNNs) trained on small patches extracted, at various magnifications, from H&E-stained WSI of rat tissue sections. The Inception-v3 network outperformed other architectures, with a patch-level test set accuracy of up to 83.4% (top-3 accuracy: 96.3%), and was therefore used for the subsequent analyses.

### Analysis of neural network embeddings

The penultimate layer of the Inception-v3 network can be considered an “embedding” of an input image patch in a high-dimensional vector space. We collected the embedding vectors for all patches from the rat test set and used nonlinear dimensionality reduction in order to visualize and interpret tissue topology in the two-dimensional plane. Patches from the same tissue formed one or more groups, often distinct from other tissues. Overlaps between groups often corresponded to similar histological structures present in more than one tissue. More in-depth exploration of the embeddings revealed that the network had learned a representation for histologically relevant substructures that had neither been annotated nor trained for.

### Cross-species predictions and transfer learning

Using control animals from studies in minipig and non-human primates (NHP), we first show that the rat histology model is able to recognize some tissues in other species, but that overall tissue recognition does not generalize well across species. However, we then demonstrate that the rat histology model is a better starting point for new tasks in the histology domain than a generic ImageNet model. In particular, we show that it leads to improved accuracy with reduced amounts of required training data when developing a new model of NHP histology. We suggest that similar observations will hold true for other machine learning tasks in a histopathological context, for which our model might provide a suitable starting point.

## Results

### Tissue recognition

Our primary objective was to assess the predictive performance of three standard neural network architectures, VGG-16 (Simonyan and Zisserman, 2015), Inception-v3 (Szegedy et al., 2016), and ResNet-50 (He et al., 2016), at recognizing a comprehensive catalogue of 46 tissues at different magnifications. In addition, to assess how the models perform in complex situations, we have included samples corresponding either to anatomical morphological variations in the brain or along the gastrointestinal tract, or complex tissues such as the eye.

The accuracy of tissue recognition was assessed using manually curated and annotated whole slide images (WSIs) from 46 normal rat tissues (Table 1) collected in six independent preclinical toxicology studies and split into training (4 studies; 1,183 slides) and test (2 studies; 507 slides) sets. Separate models were trained from patches extracted from the outlined tissues at different magnifications ranging from the scanned resolution (0.252 microns per pixel or mpp) to a 32-fold downsampling (8.064 mpp). As the pixel size of the patches was kept constant across magnifications, the actual area sampled from the slide varied from 0.003 to 3.26 mm^2^, and hence also the amount of contextual information available on a patch. For each magnification, VGG-16, Inception-v3 and ResNet-50 networks – initialized on ImageNet – were trained on patches generated at the given magnification. During training, each patch was subjected to a random rotation and a random staining modulation according to the method described in (Tellez et al., 2018). This sample augmentation aims to render the classifiers robust to changes in orientation and variations in staining.

**Table 1.**
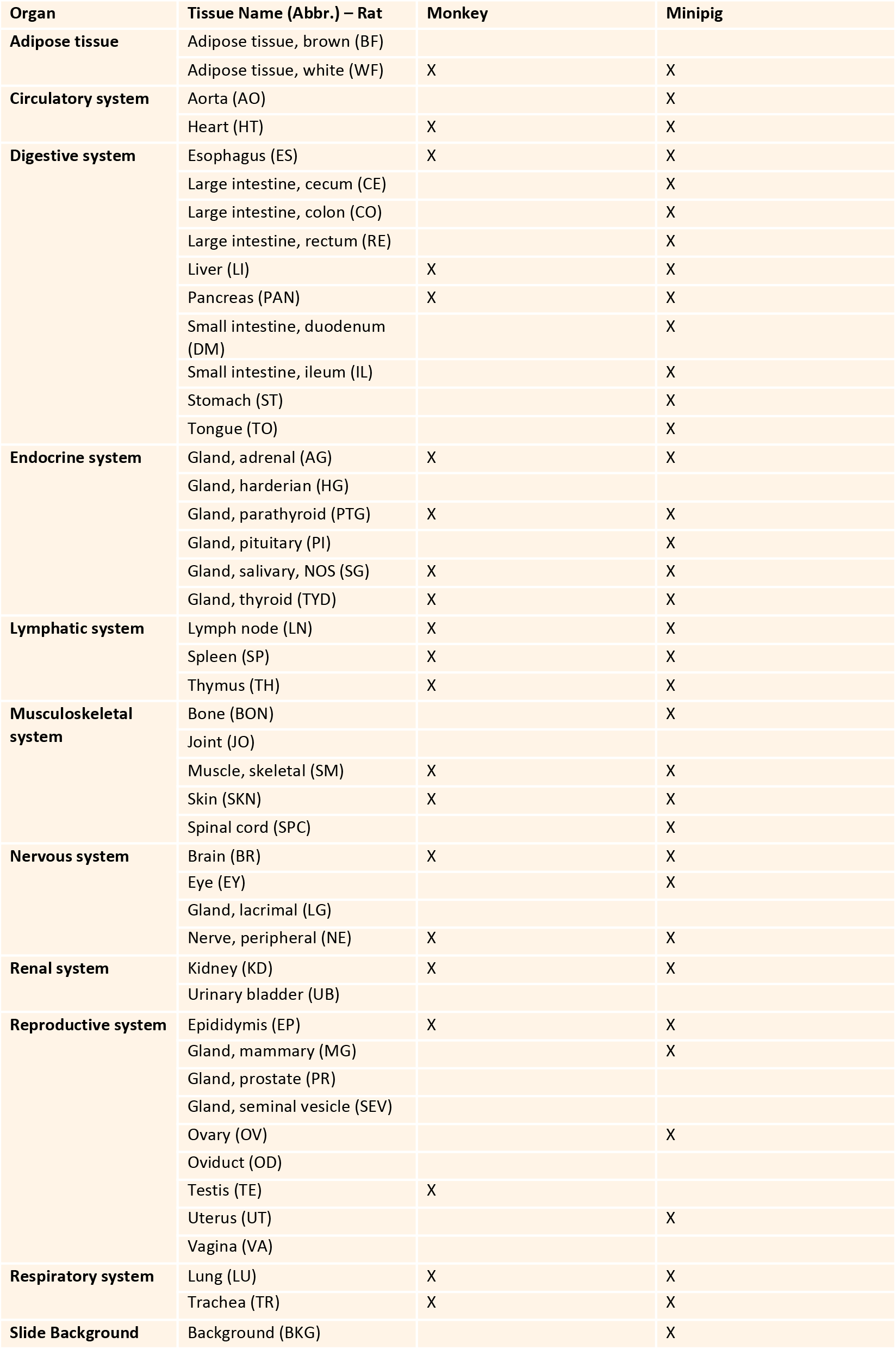
The rat tissue catalog (N=46), consisting of 45 tissues and one non-tissue class, BKG, to predict the slide background, grouped by organ system. Additional columns indicate which of these classes were available for the other species: NHP (N=21), minipig (N=36).

The accuracy of patch classification ranged from 55.1 to 83.4%, depending on magnification and architecture (Figure 1A). It was higher at low magnification, likely due to the higher amount of context available at higher mpp levels. At all magnification levels, Inception-v3 outperformed the other network architectures. The performance gap to VGG-16 was especially large, with 74.9% patch-accuracy at 8.064 mpp compared to 81.4 and 83.4% for ResNet-50 and Inception-v3, respectively. Therefore, the Inception-v3-based models were chosen as the basis for all subsequent analyses.

**Figure 1.**
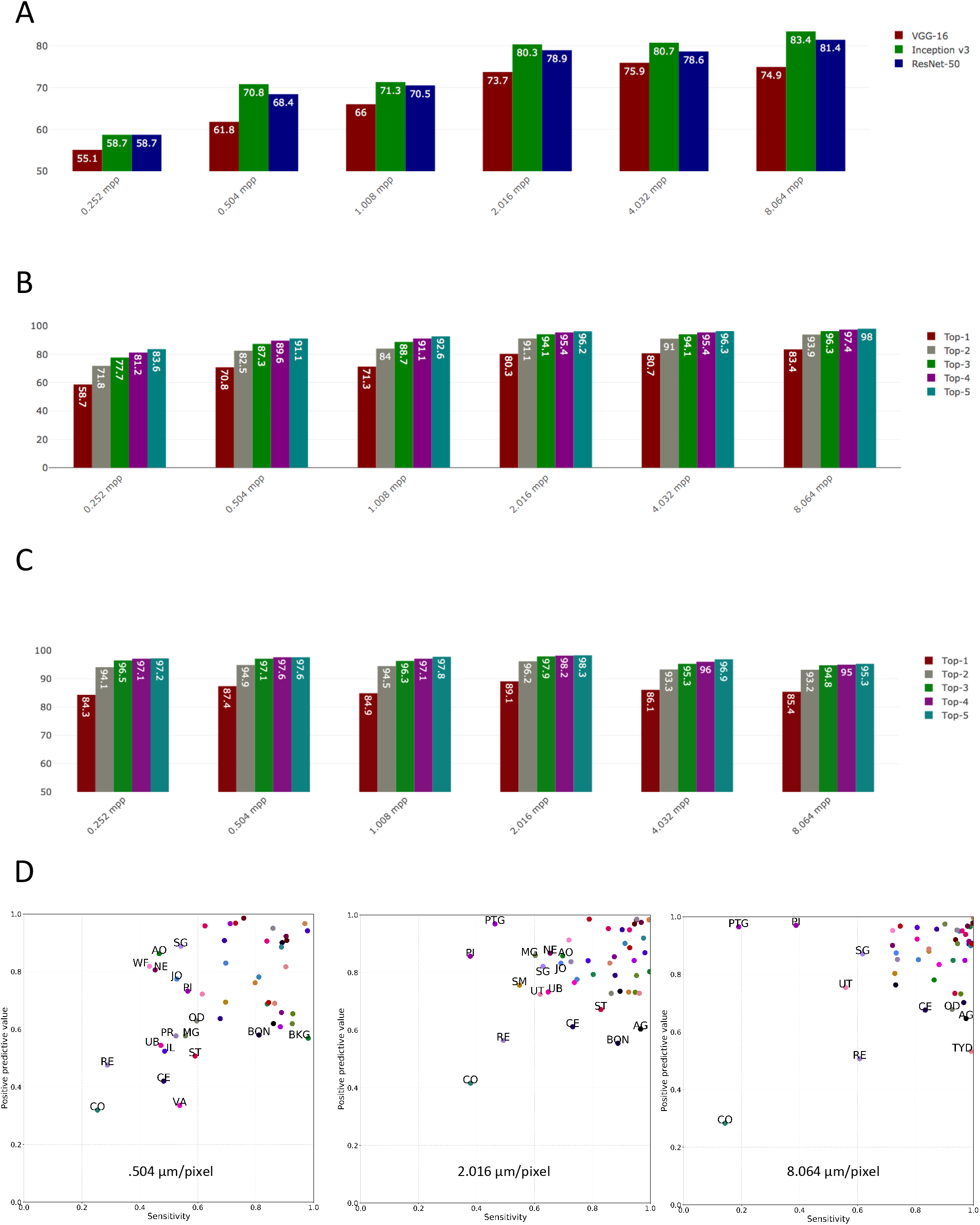
Accuracy of prediction by CNN architecture and magnification level (A). Accuracy of the top-5 predictions at the patch level by magnification, using the Inception-v3 network (B). Accuracy of the top-5 predictions at the region level by magnification, using the Inception-v3 network (C). Positive predictive value and sensitivity of the Inception-v3-based models at 0.504, 2.016, and 8.064 mpp, for each tissue (D). Tissue class abbreviations (labels are shown only below a threshold of positive predictive value and sensitivity: 0.6 was used for 0.504 mpp, and 0.7 for 2.016 and 8.064 mpp): AO: aorta, SG: salivary gland, WF: white fat, NE: nerve, JO: joint, PI: pituitary gland, OD: oviduct, BON: bone, BKG: background, PR: prostate, MG: mammary gland, UB: urinary bladder, IL: ileum, ST: stomach, RE: rectum, CE: caecum, CO: colon, VA: vagina, UT: uterus, PTG: parathyroid gland, SM: skeletal muscle, AG: adrenal gland, TYD: thyroid gland.

Given that the models were intentionally trained to differentiate morphologically similar tissues such as the different segments of the intestinal tract, the absolute accuracy of tissue prediction may not reflect fully the learning of histologically relevant high-level features. To explore this concept, the top-N accuracy was measured for models based on the Inception-v3 architecture. This metric considers a class prediction as correct if it is among the top N predictions (ranked by the probability assigned to the tissue classes by the network). At 8.064 mpp, these accuracies are 83.4% for top-1 (as reported above), 93.9% for top-2 and 96.3% for top-3 (Figure 1B). The performance gap between top-1 and top-2 was particularly large (10.5 percent points) but subsided between top-2 and −3 (2.4 percent points) and even lower for higher N.

Beyond patch classification, we also investigated test set predictive performance at the region level by combining the predictions for all patches extracted from any given test set tissue outline using a majority vote (Figure 1C). This region-level test set accuracy was much higher than patch-level accuracy for 0.252 and 0.504 mpp, and it generally appeared almost constant across magnifications. Only at 8.064 mpp was the region-level accuracy exceeded by patch-level accuracy.

Patch-level sensitivity and positive predictive value (PPV) for individual tissues showed striking differences varying with magnification (Figure 1D). Unsurprisingly, the prediction of morphologically similar regions of the large intestine such as cecum (CE), colon (CO) and rectum (RE) showed low PPV and sensitivity at any magnification. The predictions for the femoro-tibial joint (JO) and bone (BON) had low PPV and sensitivity at 0.504 and 2.016 mpp but not at 8.064, probably as patches at this magnification provide a larger context with inclusion of bone and sternum. Interestingly, the predictions for parathyroid (PTG) had high PPV and sensitivity at high magnification (0.504 mpp) but while PPV remained high, the sensitivity dropped at 2.016 and further at 8.064 mpp.

Confusion happened more often among histologically related types of tissues, where morphologies were shared at higher magnification (Supplementary Figures 1–3). Not surprisingly, regions of the large intestine (CE, CO, RE) and the small intestine (DM, JE, IL) were often confused with each other, either as a single group (0.504 mpp), or as two groups corresponding to the mid- and hindgut (2.032 mpp). As shown in Supplementary Figures 1–3, similar confusions occurred between spinal cord (SPC), brain (BR) and nerve (NE), or joint (JO), bone (BON) and skeletal muscle (SM), or between thyroid (TYD) and parathyroid gland (PTG).

### Analysis of neural network embeddings

The embedding layer of the Inception-v3 network was explored through dimensionality reduction using Uniform Manifold Approximation and Projection (UMAP; McInnes et al., 2018). The UMAP projections were calculated exclusively using the embedding vector; the information about the true tissue class of the patch was only used for coloring the plot.

A distinct projection of the embedding vectors for all test set patches was generated for each magnification as the corresponding models were trained independently (Figures 2 A, D, G, J). Projections at all magnifications revealed groups corresponding to a large extent to true tissue classes. Groups were better separated at lower magnification (i.e. higher mpp). Projections from models trained at high magnification showed a central aggregate of unresolvable patches, the size of which decreased with increasing mpp. Starting from 1.008 mpp, adjacent groups corresponded to tissue classes with similar morphology.

**Figure 2.**
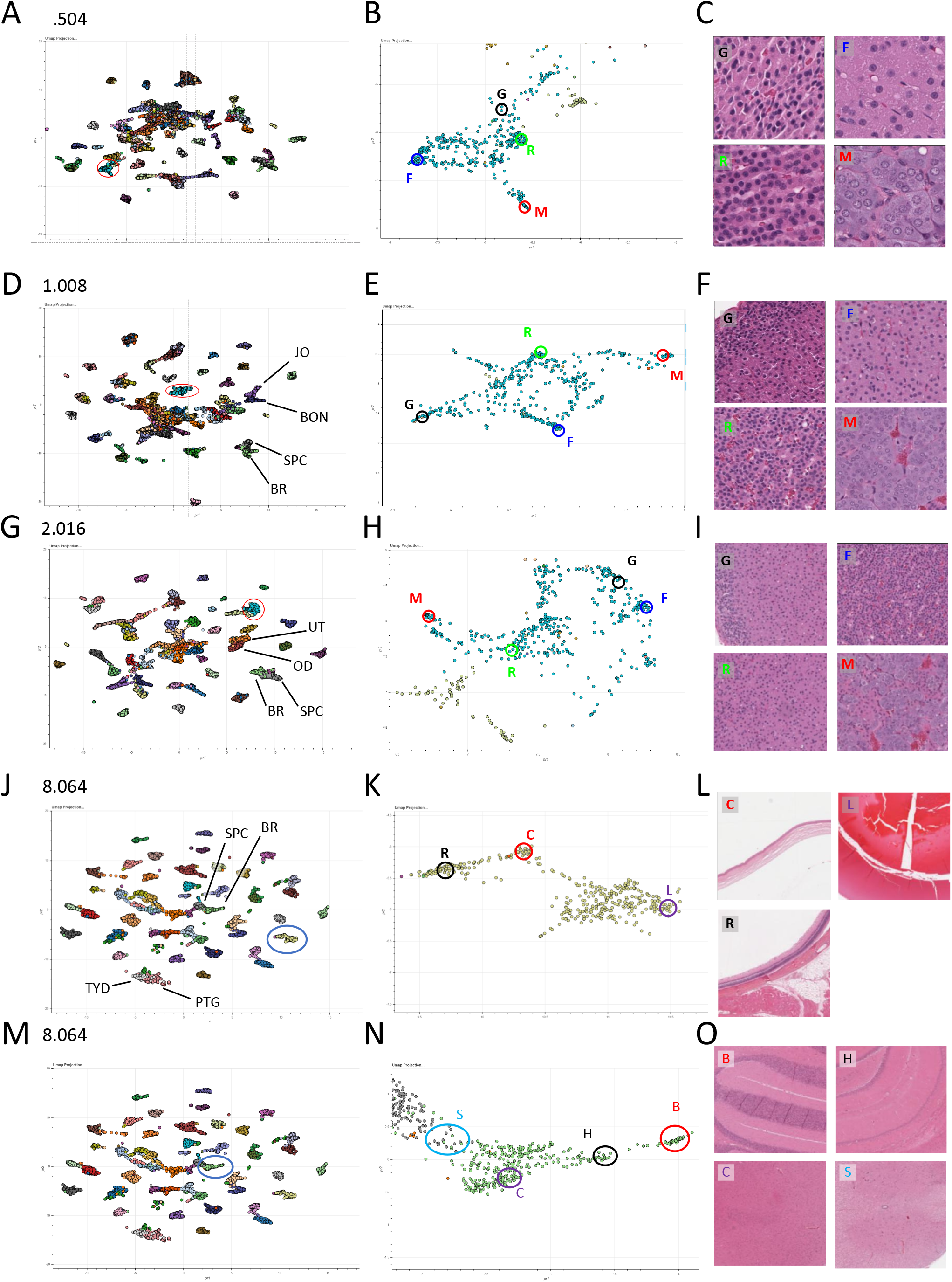
UMAP plots (left hand column) of all rat test set tissues at increasing mpp (A 0.504, D 1.008, G 2.016, J 8.064 and M 8.064), with a specific tissue cluster (circled) further represented in the clusters of the central column (B, E and H = adrenal gland, K = eye and N = brain and spinal cord). Sub-anatomical sites are represented in each central column cluster and in the respective histological images of the right hand column (adrenal gland: G = zona glomerulosa, F = zona fasciculata, R = zona reticularis, M = medulla; eye: R = retina, C = cornea, L = lens; brain and spinal cord: B = cerebellum, H = hippocampus, C = cerebral cortex, S = spinal cord).

Many clusters were not compact when given a closer inspection, and showed subclusters variably interconnected depending on the magnification at which the embeddings were generated (Figures 2 B, E, H, K, N). These subclusters corresponded to histologically meaningful regions of the tissue. For example, patches corresponding to the adrenal gland (AG) were clustered in subgroups showing different morphologies (Figures 2 B, E, H). Further exploration of the morphology of these patches showed that they correspond to distinct regions of the adrenal gland, namely, the medulla and the cortex, and within the cortex to the zonae glomerulosa, fasciculata, and reticularis (Figures 2 C, F, I). Similar histologically meaningful clustering was observed for the eye, where patches corresponding to the retina, the cornea and the lens were clearly separated from each other (Figures 2 K, L), and for the brain (Figures 2 N, O), with separation of the cerebellum, hippocampus, cerebral cortex and spinal cord.

The analysis of the embeddings also shed light onto magnification-dependent confusion between thyroid (TYD) and parathyroid (PTG), which are contiguous organs on histological sections. While at 0.504 mpp, patches from TYD and PTG were well separated (Figure 3A), at 8.064 mpp, TYP and PTG overlapped (Figure 3C). At this magnification, the parathyroid gland represents only a small part of the patch. Moreover, some thyroid or parathyroid patches containing skeletal muscle at low magnification clustered away from the glandular patches, closer to the skeletal muscle tissue group.

**Figure 3.**
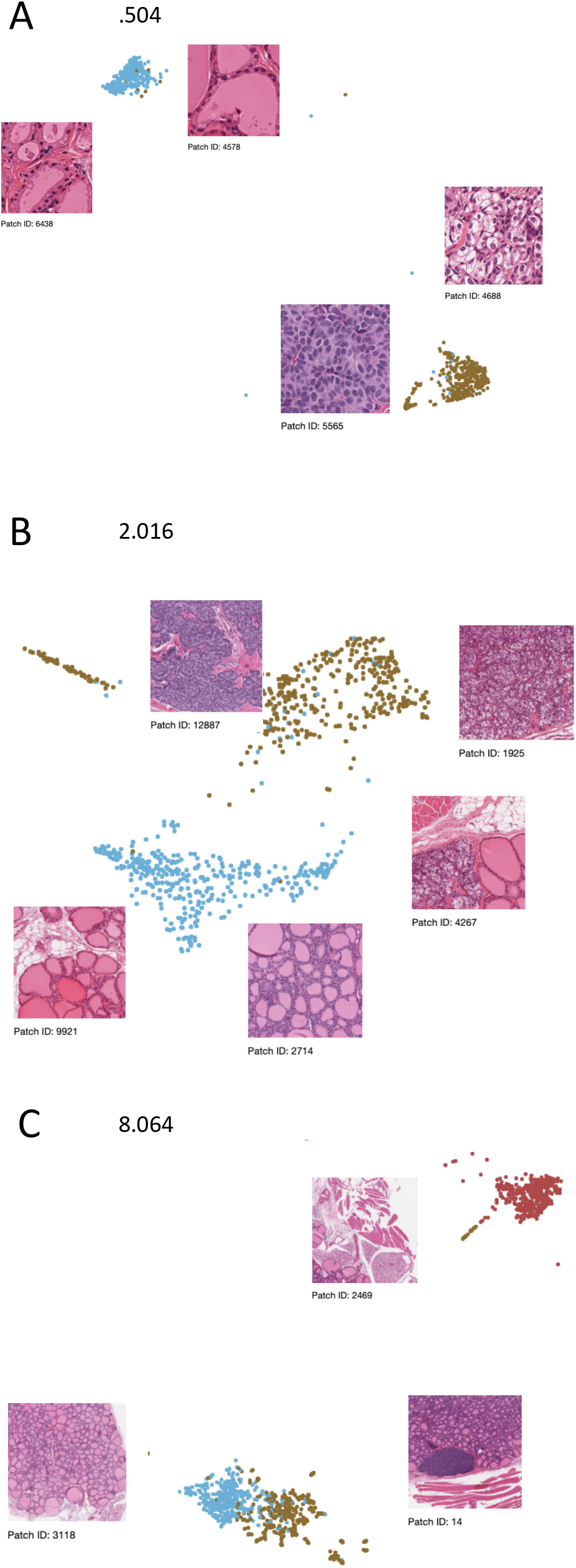
UMAP plots and respective histological images of thyroid and parathyroid gland at increasing mpp levels (light blue: thyroid, mustard: parathyroid, red: skeletal muscle).

### Cross-species predictions and transfer learning

To assess generalizability, the Inception-v3 based rat model was then used – without any retraining – for cross-species prediction of novel datasets obtained from NHP and minipig which covered 21 (NHP) and 36 (minipig) of the 46 tissues from the rat data set. At 0.504 mpp, patch-level accuracy of the predictions was 34.6% (NHP) and 39.3% (minipig). At 2.016 mpp, accuracy increased to 39.3% (NHP) and 45.2% (minipig). Finally, at 8.064 mpp, accuracy was 40.5% (NHP) and 42.0% (minipig). Many misclassifications corresponded to predicted tissues that were absent from the NHP or minipig datasets (Supplementary Figures 4 and 5). Aside from these, confusions among histologically similar tissues, as observed with the rat data, were also present. Some tissues – brain, liver, kidney, thymus, spleen, lymph node and thyroid gland – were recognizable at high accuracy in both NHP and minipig using the rat model, while other tissues – oesophagus, white fat, and tongue – were recognized well in only one species.

Given this relatively poor performance at cross-species classification without retraining, we next explored a scenario that allowed for retraining in order to adapt to a novel species. In particular, our goal was to assess whether the rat histology model would be a better starting point for species adaptation than a generic ImageNet model. To this end, another independent study consisting of WSIs from 12 NHPs was used for training, while the NHP test set was the same as in the previous cross-species prediction without retraining. Two Inception-v3 networks were trained for each magnification: one initialized with ImageNet weights, and one with the rat model weights. In addition, to determine the amount of training data necessary to achieve reasonable accuracy, a series of smaller training sets (using slides from 1, 2, 3, and 4 NHPs, respectively) was constructed from the full NHP training set of 12 animals.

Patch-level accuracies (Figure 4) were 85.8% (0.504 mpp), 92.4% (2.016 mpp), and 93.9% (8.064 mpp) when using the full training dataset (N=12 NHPs). These accuracies were substantially higher than those observed on the rat dataset, as only 21 tissue classes were available for NHP, compared to the 46 tissue classes of the rat model, with some of the most difficult tissue classes missing.

**Figure 4.**
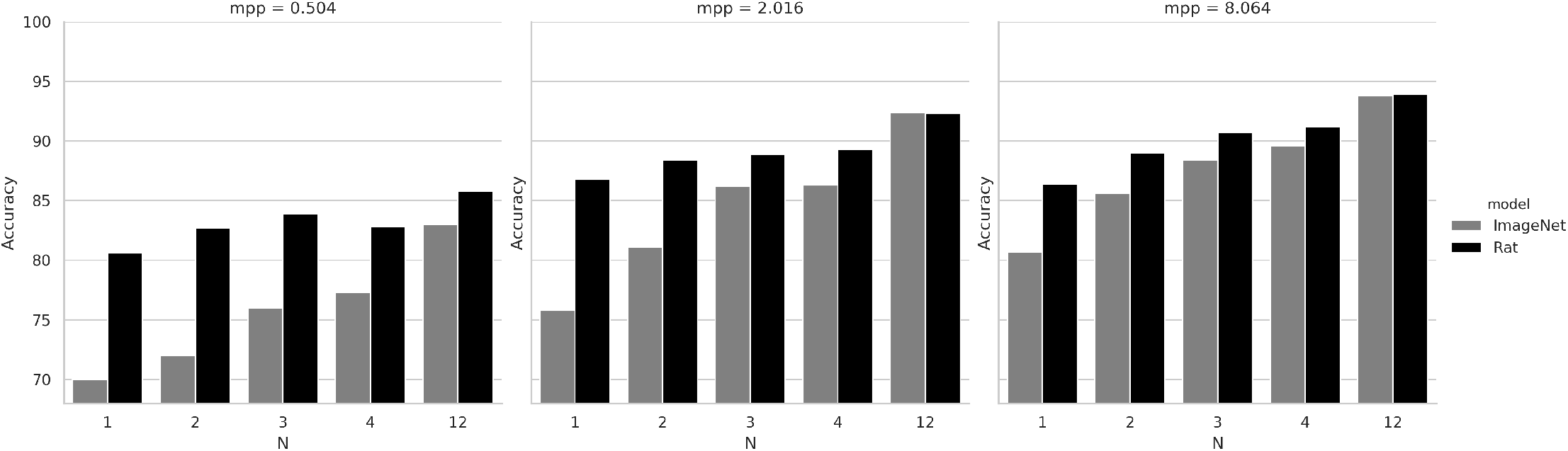
NHP test set accuracy of generic (ImageNet-pretrained) vs. domain-specific (rat-pretrained) models, after training on N=1, 2, 3, 4, and 12 (max. available) NHPs, at 0.504, 2.016, and 8.064 mpp.

Importantly, as shown in Figure 4, when only limited NHP training data (1, 2, 3, or 4 NHPs) was made available, accuracies obtained by the NHP models initialized with the weights learned from the rat dataset were consistently higher than those obtained with the standard initialization via ImageNet. For instance, the rat-initialized model using samples from 2 NHPs attained an accuracy of 82.7% at 0.504 mpp, whereas this same accuracy was only reached for the ImageNet-initialized model when using the full dataset of 12 NHPs (accuracy 83%). Likewise, at 2.016 mpp the ImageNet-initialized network required samples from 4 NHPs in order to achieve the same performance as the rat-initialized network with 1 NHP (86.3% versus 86.8%, respectively). The benefit of the rat-initialized models was less pronounced at 8.064 mpp.

## Discussion

### Tissue recognition

We have shown that standard convolutional neural network architectures can be trained to recognize a comprehensive set of rat tissues at high accuracy across a broad range of magnifications. This even applied to some extent to subtle variations of histological structure, such as segments of the small (duodenum, jejunum and ileum) and large (caecum, colon and rectum) intestine, as well as to complex organs made of several tissue types (e.g. eye or joint), for which small patches extracted at high magnification are typically difficult to classify by a pathologist in the absence of larger context. This suggests that a whole tissue model could retain some specificity towards subtle histological features.

In general, tissue prediction was increasingly reliable with decreasing magnification. This can be easily understood given that a 224 × 224 pixel patch exhibits only minimal context at high magnification (56 x 56 microns at 0.252 mpp). With decreasing magnification, more structures are present on the patches, thus facilitating tissue recognition. However, small tissues such as the parathyroid gland represented an exception. While it is recognized with consistently high PPV (meaning that if it is predicted, the prediction is usually correct) at all magnifications, the sensitivity decreased with decreasing magnification. In this case, increasing the context is detrimental to the network’s recognition as other tissues occupy increasing amounts of space on a patch (see Figure 3C for a visual example).

The majority of misclassifications were not due to unsystematic noise, but rather a result of confusions among histologically similar tissues. This was particularly prominent among segments of the large and small intestine, but also within the groups nerve, brain, spinal cord, and joint, bone, skeletal muscle, respectively (Supplementary Figures 1-3). For the latter group, the confusion is likely due to joint as a tissue being subject to interpretation, with variable amounts of bone and skeletal muscle included in the manual outlines from which the patches are generated. Misclassification also occurred based on local histological similarity in some areas of the tissue regions such as between the periarteriolar lymphoid sheaths of the spleen, and the cortex of lymph nodes (not shown). Finally, the confusion between the contiguous thyroid and parathyroid glands illustrated the role of magnification as discussed above.

For practical tissue type classification, prediction on the level of large tissue regions would be more relevant than on individual patches. The performance at the region level across magnifications was counter-intuitively constant compared to the patch-level. This may be due to predictions for patches with low mpp being more variable – and thus a majority vote being more impactful – than for patches with high mpp, which would be in line with well-known results from ensemble learning (Krogh, A. and Vedelsby, J., 1995). Our majority voting scheme can be considered a simple case of such an ensemble classifier.

### Data considerations

The reason for basing our approach on normal animal tissues as opposed to human clinical samples is tissue quality. In order to build a comprehensive atlas of normal human histology, specimens would have to be taken at autopsy, when their histological quality is usually poor due to autolysis. Notwithstanding dedicated efforts such as the GTEx tissue collection, autolysis is still an impediment, with several tissues (e.g. pancreas, stomach, prostate) having a median grade of autolysis of 2.0 (data not shown). On the contrary, large collections of tissues of high histological quality are easier to collect at the necropsy of laboratory animals used for toxicology studies. Moreover, unlike clinical surgical specimens often resected at an advanced stage of disease, a large proportion of samples from preclinical toxicology studies contain elementary lesions at various stages in relatively pure forms. This material offers a unique opportunity for the development of holistic histology and histopathology models.

A second data consideration is related to the substantial risks of overfitting or inadvertently introducing an undesired bias via the composition of the training set: In earlier stages of this work, due to limited amounts of available data, the split into training and test sets had been performed on the level of animals rather than studies. Under these conditions, test set accuracies were substantially higher than those reported here. However, as we obtained data from additional rat studies, we found that the predictive performance on these new datasets was lower. This suggested that while stratified randomization on the animal level had prevented the network from overfitting to individual animals, it instead overfitted on the study level. In order to mitigate this effect, more studies were collected, and training and test sets were split by study.

### Analysis of neural network embeddings

A popular misconception about neural networks is that they are “black box” models, i.e. harder to interpret than other machine learning methods. However, as the nature of the learned weights and the neuron activations of CNNs are eminently visual concepts, a wide variety of methods for interpreting them have been suggested. In particular, the embedding vector, i.e. the activations of the penultimate layer of a CNN can be considered a mapping from a visual into a semantic space. Proximity in this space may be thought of as semantic, rather than purely visual similarity.

As shown above, the UMAP projections at all magnifications revealed clusters that correspond to a large extent to the true tissue classes. Adjacent or partially overlapping clusters often corresponded to tissues of similar morphology. Studying the UMAP projections also aided in the interpretation of the apparent magnification-dependent confusion between thyroid and parathyroid.

Most strikingly, even though the annotations used for training the model only corresponded to coarse tissue/organ classes, the embedding layer learned by the network exhibited a degree of granularity that allowed for localizing histological substructures which had found a distinct representation without prior annotation. This suggests that our model of normal histology could be applicable beyond the task it was defined for, such as in lesion detection or in content-based image retrieval.

### Cross-species predictions and transfer learning

When applied to tissues from other species, the Inception-v3 models trained on rat showed high sensitivity and PPV only for some tissues. At the level of the individual tissues, it is interesting to note that those with very conserved structure such as liver, kidney, brain, thyroid and the lymphoid organs were classified with high accuracy across species. Not surprisingly, the gastrointestinal tract was often misclassified, which is not different from the prediction made on rat tissues.

Despite high predictive performance for some tissues, the rat model did not show sufficient overall accuracy on the cross-species prediction task to be immediately applicable. However, it provided a good basis for transfer learning to a novel species (NHP), superior to a model that had been initialized with ImageNet weights. The rat-initialized model showed its biggest benefits when training data was scarce or the training task was hard (i.e. at high magnification levels). Of note, some of the tissues for which prediction was poor in the rat dataset where not part of the NHP dataset, which – along with the lower number of classes to predict (21 in NHP vs. 46 in rat) – can explain the substantially higher accuracies seen in NHP, as compared to the original rat data.

In addition to being beneficial in the close-at-hand task of cross-species tissue classification, this approach of transfer learning from a domain-specific model may also be a suitable starting point when training for other tasks in the histology domain, e.g. the recognition of a specific type of lesion for which only limited training data is available.

### Related work

Our suggestion of reusing domain-specific models relates to the notion of pretext tasks in machine learning (Noroozi and Favaro, 2016; Noroozi et al., 2017), in which a task is learned not (only) for its own sake, but to learn a feature representation which can be useful for other tasks in the same domain. Below, we discuss a few other topics that are relevant in the context of our work.

During the writing of this article, another study appeared which also aimed at predicting tissue types from patches, and with a similar approach to ours, however with a focus on a software framework for reproducible machine learning (Bizzego et al., 2019). While this effort is in parts overlapping with our work, there are important differences between the two efforts. Our work focuses on several animal models (rat, NHP, minipig) instead of the human GTEx dataset alone. For each animal model, we collected several independent studies and ensured that our training and test sets come from separate studies. As was discussed in detail above, these are crucially important features required for obtaining robust models and reliable (non-inflated) performance estimates. In addition, we evaluate the effect of different magnification levels on the models. On the deep learning side, we train all our models end-to-end, including the convolutional feature generation layers and we evaluate transfer learning between species. In the remaining paragraphs, we discuss further lines of related work.

#### Multi-scale CNNs

The scanned slide images are of such high resolution that they contain diverse structural and textural information on different levels of magnification, from the level of individual nuclei to the level of entire tissues. Instead of training a separate network for each level of magnification, as we have done in this work, an obvious extension would be to build a convolutional network which accepts such hierarchical input, i.e. patches at several levels of magnification. This could potentially combine the benefits of having enough context on a patch with the advantages of lower-magnification patches in recognizing smaller tissue structures, as was observed in an application of multi-scale CNNs to high-content cellular images (Godinez et al., 2017).

#### Visual explanations

In terms of interpreting the representations learned by the network, besides analyzing the embedding layer by means of nonlinear dimensionality reduction, a great number of approaches has been suggested that focus on different aspects of the model. One approach that will likely yield to greatly improved insight into what kind of structures or textural features are most relevant for a given class prediction is the Grad-CAM method (Selvaraju et al., 2016). Initial experiments with this technique for deriving “visual explanations” have been promising (data not shown).

#### Semantic segmentation; weakly supervised approaches

Besides our patch-based approach, several other machine learning paradigms have been applied to histopathology images. The most relevant of these are semantic segmentation using fully convolutional neural networks (rewieved in Wang et al., 2019) and weakly supervised learning (multiple instance learning). The advantage of the latter approach is that unlike our approach or semantic segmentation, it does not rely on outlining structures within an image (which is often resource-intensive) but rather only requires a slide-level label. Weakly supervised learning is most applicable when large numbers of slides with reported findings are already available (e.g. Campanella et al., 2019).

### General conclusion

Our results show that it is possible to train models on the extensive diversity of tissue histology. Both the UMAP analysis and the cross-species transfer learning indicate that training on the coarse tissue annotations on one species yields embeddings that are much richer from a feature perspective than the original task would suggest. This is closely related to the concept of a pretext task, which is a useful form of reusing a learned representation when labeled data for a task of interest is scarce, but much more data is available that is either unlabeled or labeled for a different task. This opens the perspective of creating general-purpose models for the classification of lesions. Such models would reduce the need for difficult and labor-intensive manual labeling of individual lesions and allow to work from entire tissues.

Holistic histology classification models such as those described here can also be useful to generate numerical data from any histological picture or series of tiles from WSIs, thereby enabling content-based image retrieval and mining of morphological data in the histopathology domain.

## Materials & Methods

### Dataset

Hematoxylin and eosin (H&E) stained slides from independently processed preclinical toxicology studies were retrieved from archives. To establish the rat models, six studies performed in Han-Wistar rats (*Rattus norvegicus*) provided 1,690 WSIs of 46 different tissues with at least 20 slides per tissue, covering nine organ systems (Table 1; for simplicity the slide background was included as a tissue class). Organs representing subtle variation of tissue structure (such as different segments of the intestine), or complex arrangements of tissues (eye, joint) were intentionally included as extreme cases.

Four studies were selected for the training dataset (1,183 slides) and two for the test dataset (507 slides). To evaluate the cross-species translatability and species-specific models, WSIs from minipig (*Sus scrofa domesticus*) and cynomolgus NHP (*Macaca fascicularis*) were generated from the control arm of one (minipig) and two (NHP) studies, covering subsets of 36 and 21 tissues, respectively.

Whole slide images (WSIs) were generated at 40x magnification (0.252 microns per pixel — mpp) using either Aperio or Hamamatsu scanners. All WSIs were reviewed by pathologists to ensure absence of lesions. Then, tissues were outlined and labeled according to a catalog of tissue types (see Table 1). Manual outlines defined regions on the WSIs from which patches of size 224 × 224 pixels were extracted at the various magnification levels corresponding to 0.252, 0.504, 1.008, 2.016, 4.032 and 8.064 mpp. As the pixel size of patches is constant across all magnifications, low magnification (high mpp) patches cover a larger area of the underlying slide as compared to high magnification patches. For example, a patch at 0.252 mpp covers an area of 56.4 × 56.4 microns on the slide while a patch created at 8.064 mpp covers an area of 1.8 × 1.8 mm (7,168 × 7,168 pixels at the original scanning magnification).

To support the learning of tissue boundaries, we also allowed patches to overlap with the outside of a tissue: A patch is valid for a region if its center is contained within the region. This approach is suggested by empirical findings of improved predictive performance when incorporating challenging boundary patches into the training set (Janowczyk and Madabhushi, 2016).

To ensure that the training and test sets are well balanced, we aimed at having equal numbers of patches for each tissue in each dataset (approx. 30,000 and 2,000 patches/tissue in the training and test set, respectively). However, the vastly different macroscopic size of some of these tissues made it problematic to sample the same number of patches for each tissue due to oversampling concerns. For example, in the training set, 32,600 mm^2^ of brain tissue and 27,200 mm^2^ of lung tissue were available, in contrast to only 25 mm^2^ of parathyroid gland tissue (Supplementary Table 1). In order to address this issue, the following scheme was implemented:

The midpoints of the patches were picked at random from the tissue area. The patches were then generated at all mpp levels for the same patch midpoint and the same random rotation. In order to avoid oversampling for tissues with low areas, the number of patches was decreased so that the average patch density at the mpp level of 0.504 was at most 1. For the training set, 34 of 46 tissues had the full 30,000 patches, another 11 tissues between 10,000 and 30,000 patches and parathyroid gland with about 2,000 training patches. For the test set, 2,000 patches were sampled which was low enough to create the full set of patches for each tissue. For a complete list of tissue areas as well as number of sampled patches for the training and test sets see Supplementary Table 1.

### Machine learning approach

For the three network architectures adopted in this study, VGG-16, Inception-v3, and ResNet-50, we used their implementations available in keras (with tensorflow back-end). The weights trained on the ImageNet dataset were used for initialization of the convolutional layers. For all fully connected layers, we use random initialization of the weights. The final layer has one unit for each tissue class (and one unit for the additional “slide background” class) and is set to have a softmax activation function outputting a probability vector.

The RGB channels of the image patches are scaled to the interval [0,1]. The training and test set patches were created and saved not during but before the learning process and subjected to a random rotation. This was done beforehand in order to avoid having to store additional slide context which would be necessary for obtaining a novel rotated rectangular patch “on the fly”. This means that in our approach each patch is used with the same rotation during each epoch. On the other hand, the stain modulation is performed during the training process (and thus differently also for the same patch across different epochs) according to the method described in (Tellez et al., 2018). Briefly, this method is based on first transforming from RGB color space to a color space consisting of one channel each for eosin, hematoxylin, and a third channel for the remaining color variation. These channels are then modified individually by a random linear transformation with parameters sampled from uniform distributions. Finally, the modified patch is back-transformed into RGB color space. We experimented with different widths for the uniform distributions, and found that a parameter choice that doubles the variation reported in (Tellez et al., 2018) as ‘typical’ resulted in the best generalization ability, while leading to pronounced changes in the visual appearance of the patches. Note that even patches from a single slide will be oriented and stained independently from each other during the training process. Patches from the test set are not subjected to any staining modulations.

For training, we use the Adam optimizer with a learning rate reduction once learning stagnates. We start out with an initial learning rate of 10^−6^ and a reduction of the learning rate by a factor 5 after 5 epochs of no improvement. We apply early stopping after 10 epochs of no improvement. During training, we updated all weights, not just the fully connected layer weights, thereby tuning the feature extraction stack of the architectures that had been pretrained on ImageNet to be better adapted to histopathology images. The same approach was also employed for the cross-species transfer learning in which we adapted the rat model to NHP. We considered an “epoch” not a full run through the entire training data, but rather through a randomized subset of 10% of the data, in order to get more fine-grained metrics of the training progress.

The performance metrics used in this work are accuracy (the fraction of samples for which the predicted class equals the true class), error rate (1-accuracy), sensitivity for class t (the fraction of samples from class t that are correctly recognized), and positive predictive value for class t (PPV; the fraction of samples predicted as t that truly belong to class t).

For the UMAP projection of the embeddings we use the software library provided by the UMAP authors (github.com/lmcinnes/umap).

## Figure and table legends

**Supplementary Table 1:** List of annotated areas as well as number of sampled 224 × 224 pixel patches per tissue type for the rat training and test datasets.

| Tissue name | Area (mm<sup>2</sup>) |  | # of patches |  |
| --- | --- | --- | --- | --- |
|  | Training | Test | Training | Test |
| Brain | 32641.4 | 11751.6 | 30053 | 2038 |
| Lung | 27261.1 | 3295.2 | 30063 | 2007 |
| Liver | 17577.7 | 2390.9 | 30049 | 2006 |
| Kidney | 15861.8 | 11874.5 | 30039 | 2028 |
| Background | 8644.4 | 187.7 | 30337 | 2011 |
| Heart | 6305.8 | 1129.4 | 30027 | 2005 |
| Epididymis | 6012.1 | 5401.0 | 30017 | 2025 |
| Testis | 5125.4 | 3960.0 | 30015 | 2021 |
| Bone | 4878.0 | 280.1 | 30041 | 2005 |
| Eye | 4658.1 | 264.0 | 30033 | 2004 |
| Spinal cord | 4533.0 | 558.8 | 30033 | 2016 |
| Gland, salivary, NOS | 4322.6 | 348.3 | 30035 | 2012 |
| Stomach | 4006.9 | 2011.1 | 30031 | 2014 |
| Joint | 3818.5 | 504.2 | 30012 | 2002 |
| Muscle, skeletal | 2620.3 | 2229.9 | 30061 | 2032 |
| Uterus | 2567.4 | 156.9 | 30024 | 2003 |
| Tongue | 2213.2 | 5051.0 | 30029 | 2024 |
| Pancreas | 2168.2 | 626.7 | 30058 | 2011 |
| Lymph node | 1992.7 | 2167.6 | 30077 | 2030 |
| Gland, harderian | 1725.9 | 321.0 | 30030 | 2004 |
| Thymus | 1564.4 | 571.7 | 30036 | 2011 |
| Spleen | 1305.4 | 966.3 | 30040 | 2024 |
| Skin | 1224.9 | 90.7 | 30030 | 2002 |
| Urinary bladder | 1202.1 | 856.4 | 30025 | 2022 |
| Gland, mammary | 1153.4 | 39.0 | 30027 | 2003 |
| Vagina | 1080.5 | 194.7 | 30023 | 2002 |
| Gland, lacrimal | 884.8 | 3163.3 | 30027 | 2017 |
| Ovary | 837.2 | 175.6 | 30026 | 2003 |
| Gland, seminal vesicle | 700.1 | 1339.5 | 30011 | 2021 |
| Gland, prostate | 670.2 | 5825.2 | 30008 | 2017 |
| Gland, adrenal | 626.5 | 295.5 | 30035 | 2006 |
| Nerve, peripheral | 521.4 | 604.7 | 30047 | 2044 |
| Small intestine, duodenum | 435.2 | 152.5 | 30024 | 2004 |
| Gland, thyroid | 408.2 | 648.7 | 30022 | 2025 |
| Large intestine, cecum | 366.1 | 319.1 | 28735 | 2010 |
| Adipose tissue, brown | 318.5 | 69.8 | 25044 | 2026 |
| Large intestine, colon | 284.8 | 191.9 | 22357 | 2011 |
| Large intestine, rectum | 279.4 | 378.0 | 21932 | 2009 |
| Small intestine, ileum | 246.7 | 144.9 | 19370 | 2010 |
| Oviduct | 222.4 | 30.5 | 17465 | 2002 |
| Adipose tissue, white | 215.5 | 474.5 | 16935 | 2041 |
| Gland, pituitary | 185.2 | 319.4 | 14541 | 2021 |
| Trachea | 143.8 | 118.7 | 11307 | 2025 |
| Esophagus | 132.1 | 177.2 | 10380 | 2030 |
| Aorta | 129.0 | 121.2 | 10148 | 2027 |
| Gland, parathyroid | 24.5 | 30.2 | 1936 | 2024 |

**Supplementary Figure 1.**
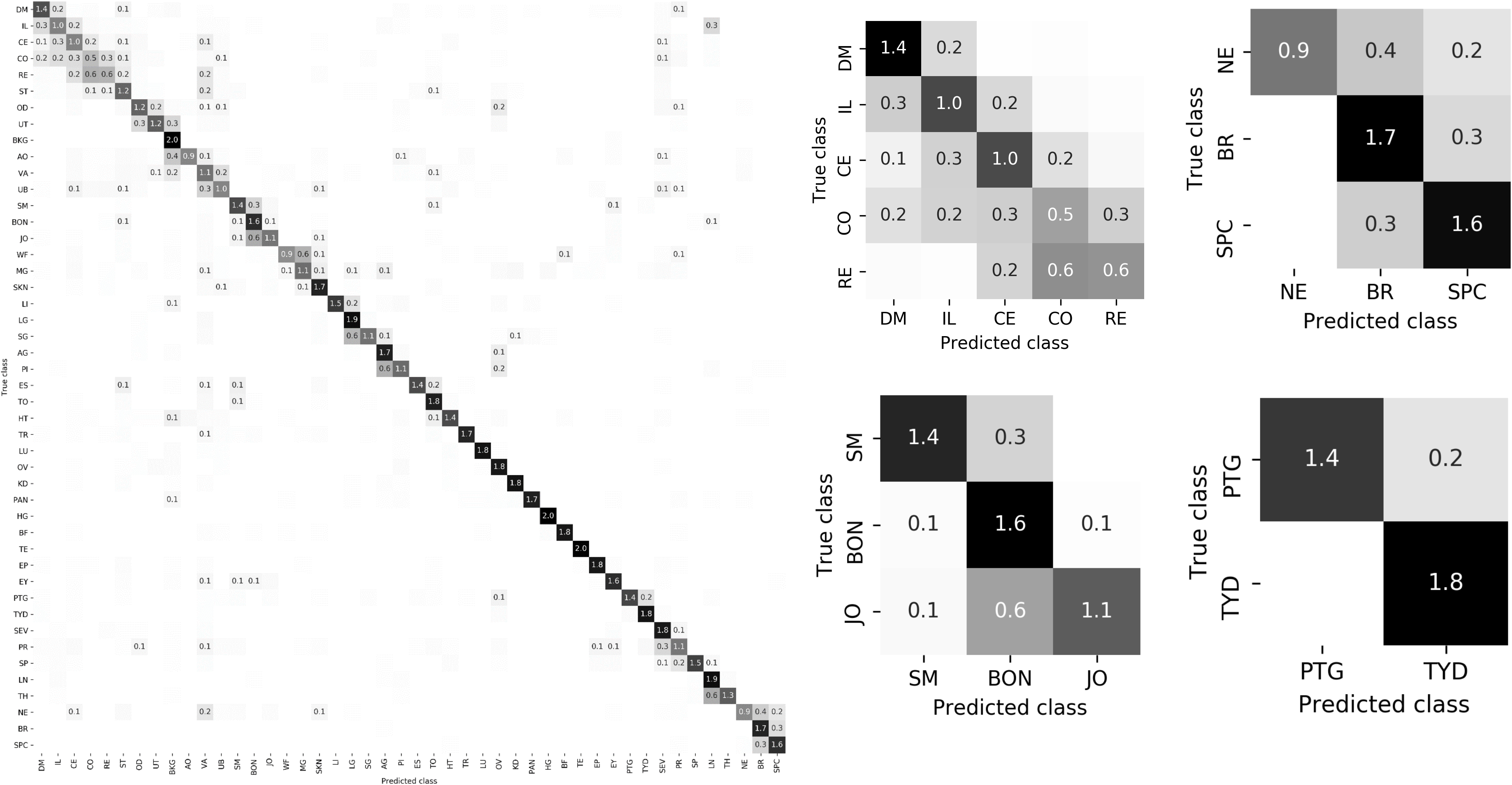
Confusion matrix for the rat model test set, 0.504 mpp, predicted using the Inception-v3 model trained on the rat training data. Prominent confusion clusters are shown enlarged on the right-hand side.

**Supplementary Figure 2.**
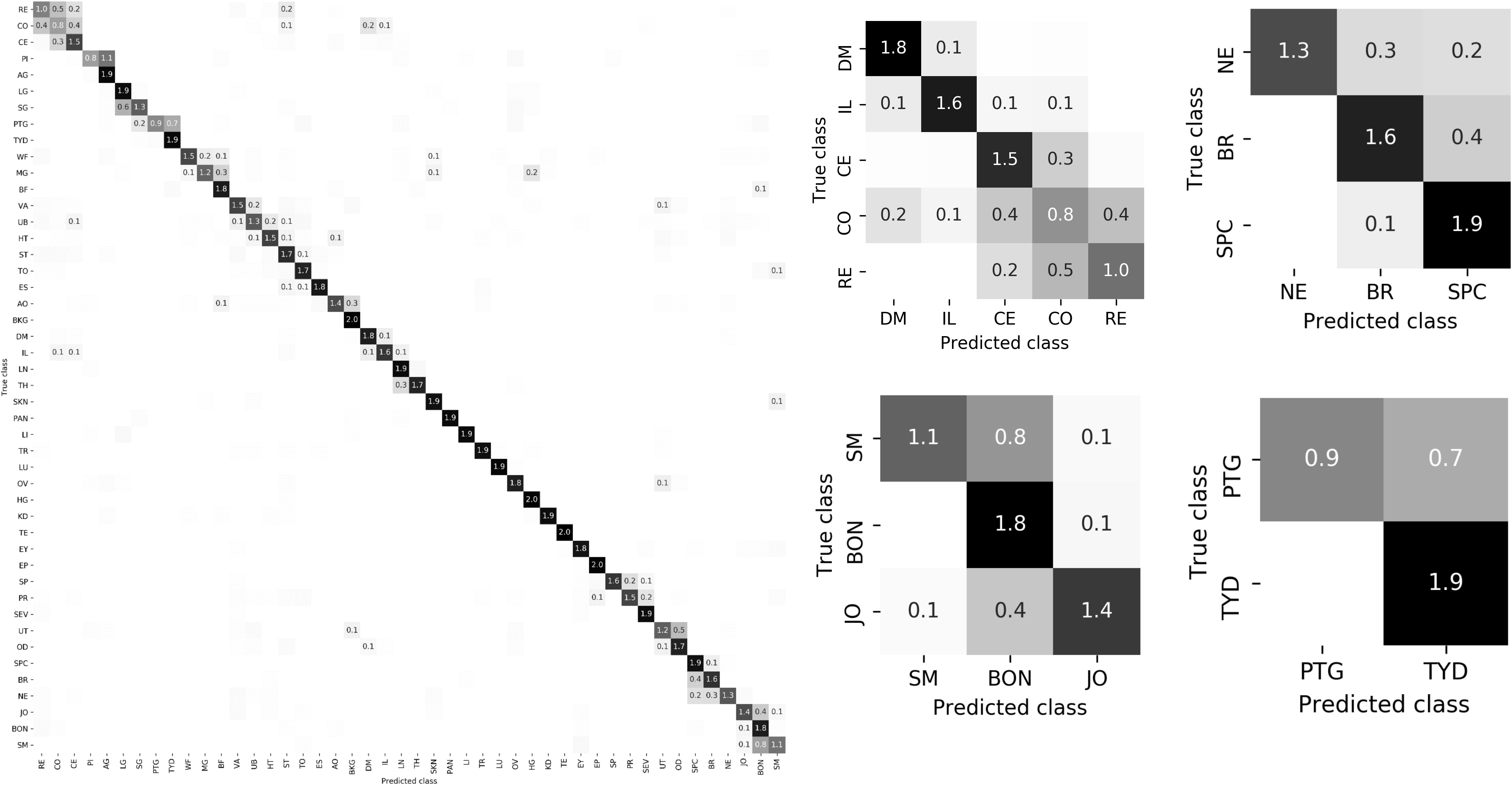
Confusion matrix rat the model test set, 2.016 mpp, predicted using the Inception-v3 model trained on the rat training data. Prominent confusion clusters are shown enlarged on the right-hand side.

**Supplementary Figure 3.**
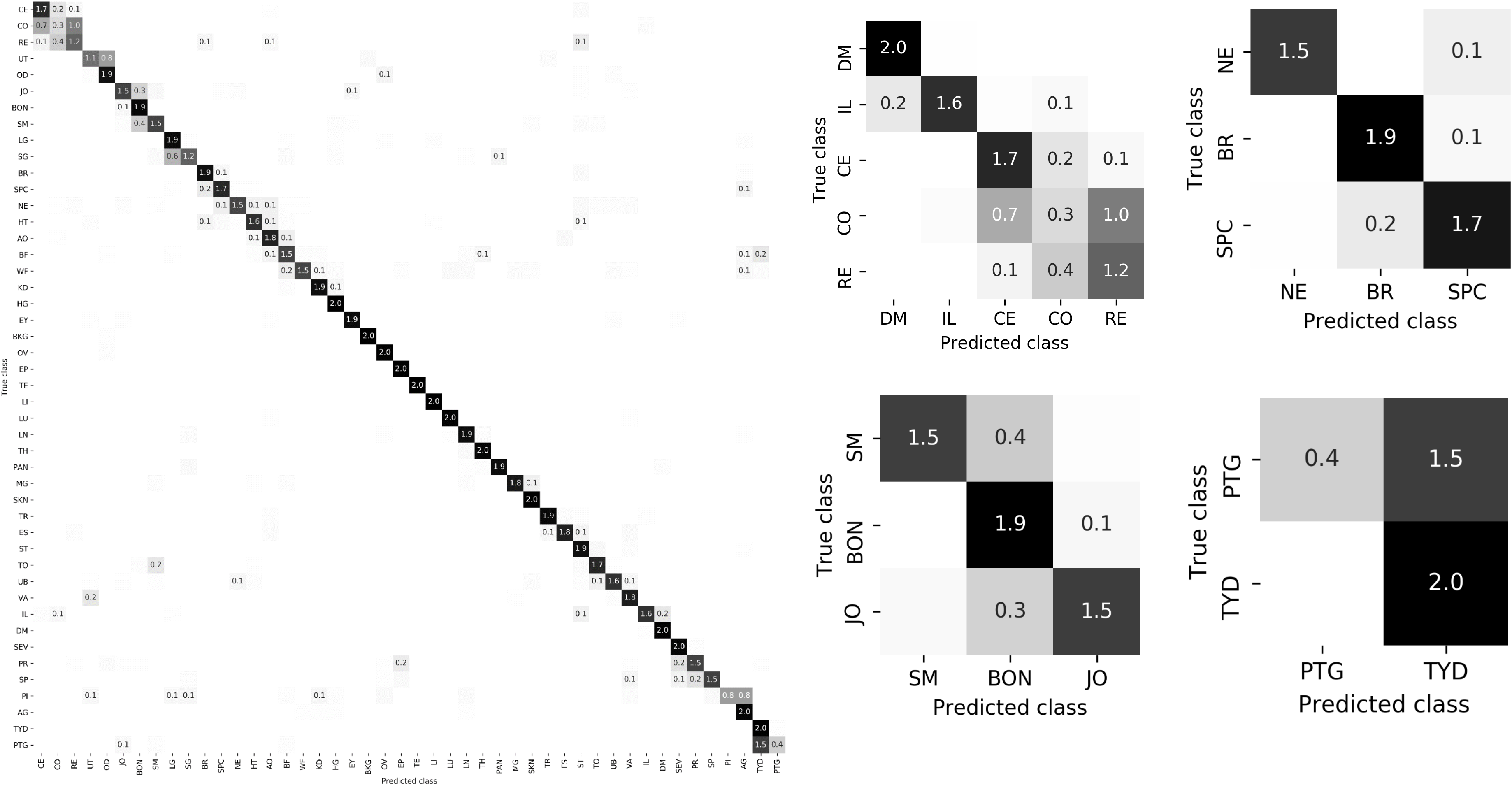
Confusion matrix rat the model test set, 8.064 mpp, predicted using the Inception-v3 model trained on the rat training data. Prominent confusion clusters are shown enlarged on the right-hand side.

**Supplementary Figure 4.**
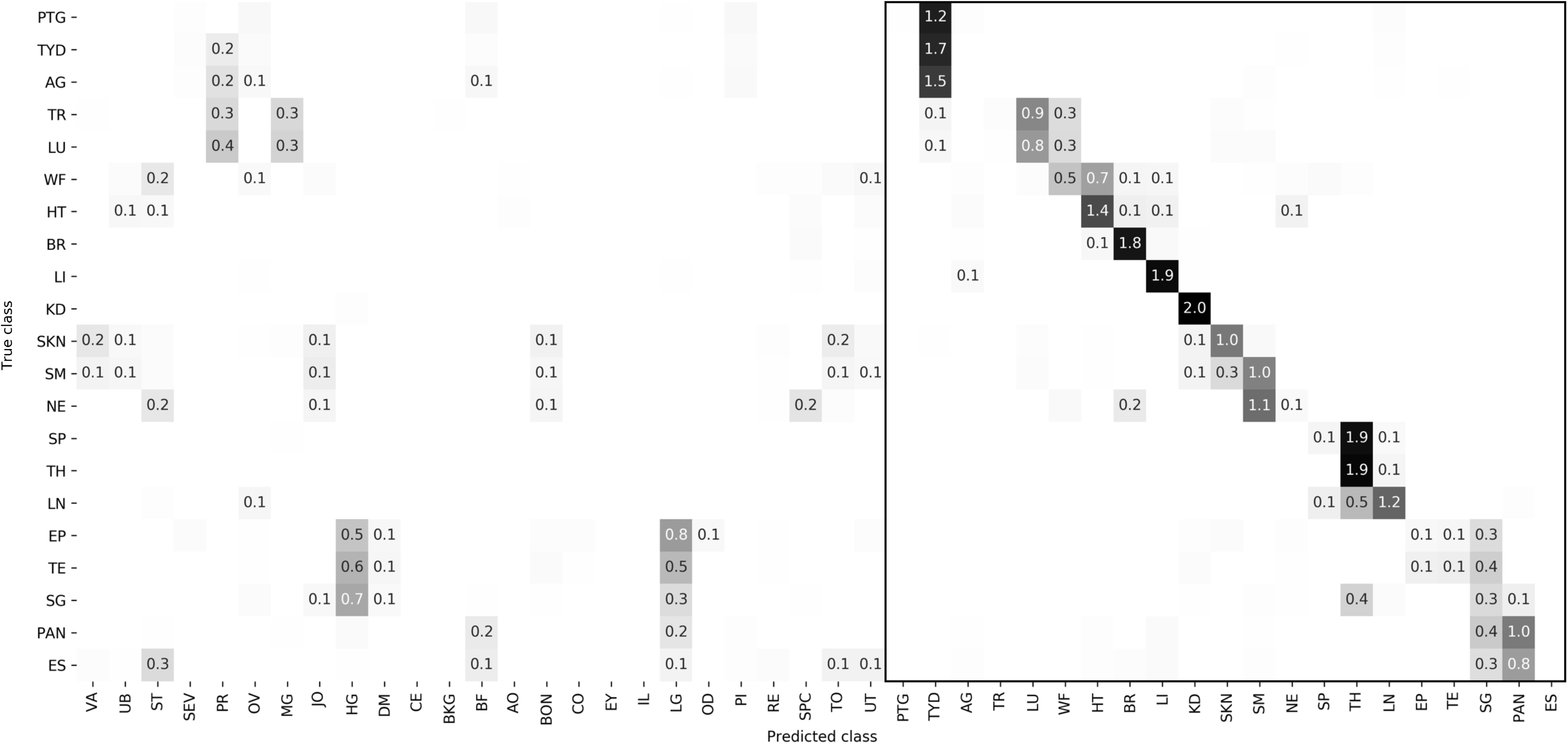
Tissue confusion matrix for the NHP test dataset at 8.064 mpp, predicted using the model trained on the rat training data.

**Supplementary Figure 5.**
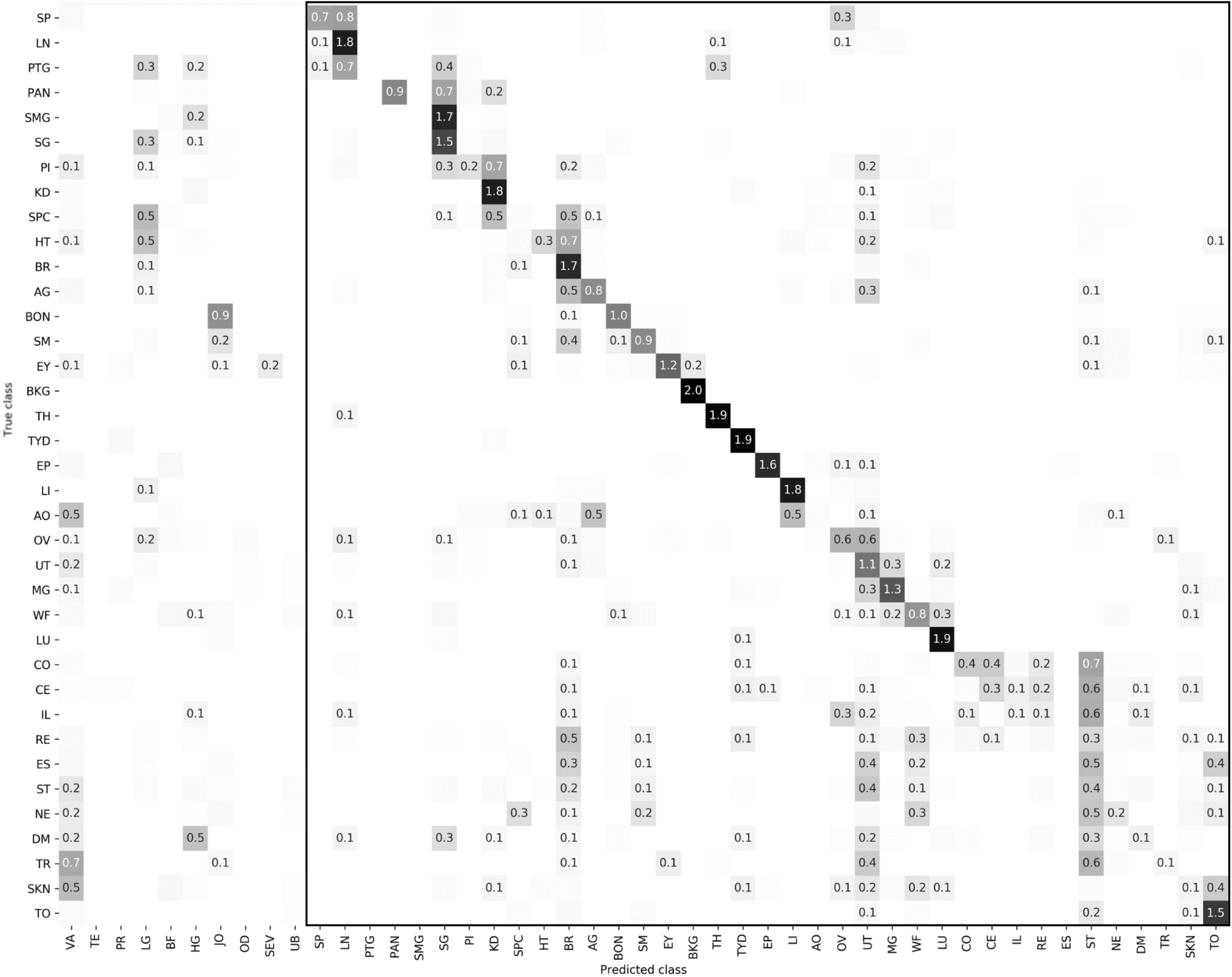
Tissue confusion matrix for the minipig test dataset at 8.064 mpp, predicted using the model trained on rat training data.

